# Thermal adaptation crosstalk with azole response through lncRNA in *Aspergillus fumigatus*

**DOI:** 10.64898/2026.03.19.713036

**Authors:** Nava Raj Poudyal, Ryan T. Mehlem, Ritu Devkota, Sourabh Dhingra

**Affiliations:** Department of Biological Sciences, Clemson University, Clemson, SC, USA 29634; Eukaryotic Pathogen Innovation Center, Clemson University, Clemson, SC, USA 29634; Department of Genetics and Biochemistry, Clemson University, Clemson, SC, USA 29634

**Keywords:** long non-coding RNAs, temperature adaptation, azole drug response, drug resistance, host-pathogen interaction

## Abstract

Fungal pathogens are adapting to increased temperatures altering host-pathogen interactions, disease patterns, and response to the antimicrobial drugs. Here, we show that thermal adaptation to 42°C leads to reversible changes in fungal colony size and azole drug response in the human pathogenic fungus *Aspergillus fumigatus*. Importantly, this adaptation is mediated by the lncRNA, *afu-182,* whose RNA levels negatively correlate with temperature. Growth at a lower temperature or ectopic upregulation of *afu-182* RNA levels reverses the temperature adaptation. Previously, we have shown that Δ*afu-182* strains produce worse disease outcomes in a murine model of invasive pulmonary aspergillosis (IPA). Here, more importantly, we show that the overexpression of *afu-182* in clinically azole-resistant isolates increased survival in a murine model of IPA. Taken together, fungal adaptation to increased temperature leads to a decrease in *afu-182* RNA levels that is associated with worse disease outcomes upon azole treatment. This provides a framework to take temperature into account when analyzing the rise in azole MIC in environmental and clinical isolates.

## Introduction

Overcoming human physiological temperature is considered to be a limiting virulence factor for pathogenic fungi (1). Consequently, increasing environmental temperatures are leading to a rise in human fungal diseases (2–6). The subsequent thermal adaptation is likely to add complexity in disease progression and treatment outcomes for diseases that are currently treatable (7, 8). *Aspergillus fumigatus* is a saprophytic fungus with cosmopolitan distribution, capable of causing invasive pulmonary aspergillosis (IPA) in immunocompromised patients and is the causative agent for most common mold associated infections worldwide (9). *A. fumigatus* can infect mammals (37°C) and birds (39-42°C) and can grow at temperatures up to 52°C while maintaining a saprophytic lifestyle (∼25°C) in the environment (10). Most of the infections are caused by azole-susceptible *A. fumigatus* (ASAF) isolates; however, infections caused by azole-resistant *A. fumigatus* (ARAF) isolates are on the rise (11, 12).

The rise in azole resistance is mostly associated with the use of agricultural azoles and fungal adaptation to continuous azole stress (13, 14). However, an increased percentage of ARAF strains are isolated from compost heaps associated with agricultural waste (15–18), suggesting a correlation between high temperature adaptation often associated with compost and azole response. Consequently, a link between temperature adaptation and azoles was recently shown in the human pathogenic fungus *Cryptococcus neoformans* (19). Additionally, *A. fumigatus* isolates from animals (especially poultry), and their close environment, have increased tolerance towards the azole antifungal drugs *in vitro,* with some isolates growing at concentrations approaching or exceeding human clinical breakpoints (20). Since most of the companion animals and domestic animals, including poultry, have body temperatures higher than that of humans (21), the role of fungal thermal adaptation and its link with azole response warrants further study.

Here, using a laboratory adaptation experiment, we show that *A*. *fumigatus* thermal adaptation at 42°C leads to changes in colony morphology, including colony size and fungal surface attachment. More importantly, thermally adapted strains grow better in sub-MIC concentrations of azoles and produce worse disease outcomes in a corticosteroid murine model of invasive pulmonary aspergillosis when treated with azole drugs. We show that this adaptation is reversible and is mediated by the lncRNA *afu-182*, which regulates fungal sub-MIC azole response *in vitro* and *in vivo* (22). Additionally, we show that overexpression of *afu-182* in clinically azole-resistant *A. fumigatus* isolates improved treatment outcomes in a murine model of IPA, indicating the role of *afu-182* in azole treatment outcomes in ARAF strains.

Taken together, our data show that *A*. *fumigatus* adaptation to increased temperature correlates with changes in *afu-182* RNA levels that ultimately affect azole treatment outcomes without changing the azole drug MIC. Previous reports show a high degree of correlation between treatment failure rates and increased MIC values for ARAF strains (23); however, here, azole treatment outcomes show a positive correlation with increasing RNA levels of *afu-182*, we suggest a better classification than susceptible/resistant isolates for *A*. *fumigatus* in the future.

## Results

### Temperature adaptation influences colony morphology and surface attachment in *A. fumigatus* but is dispensable for virulence

*Aspergillus fumigatus* is a saprophytic fungus that mainly lives in the environment (∼25°C); however, it adapts to mammalian and avian body temperatures (37°C-43°C) during infection (10). *A. fumigatus* is also constantly exposed to temperatures above 50°C in compost piles (17). To better understand the fungal adaptation at increased temperature during infection, we used an experimental adaptation approach and cultured *A. fumigatus* at 42°C, collecting spores and re-inoculating every 48 hours for 12 generations (Figure 1A, 1G^T^-12G^T^ strains, where numbers 1-12 denote the generation and T denotes the temperature variable). Temperature adaptation led to a change in qualitative fungal morphology (Figure 1B, yellow arrows). A significant increase in colony diameter was observed in the 8G^T^ (p=0.008) and 12G^T^ strains (p=0.0138) at 42°C (Figure 1C, One-Way ANOVA and Tukey’s post-hoc analysis). Additionally, the adapted strain 12G^T^ showed increased surface attachment compared to the unadapted strain (Figure 1D, p=0.0074, student’s t-test). Thus, given increased colony size and surface attachment, we tested the virulence of the adapted strain 12G^T^ compared to the unadapted strain in a corticosteroid murine model of invasive pulmonary aspergillosis. However, we did not see a change in mortality (Figure 1E). Thus, temperature adaptation increased fungal fitness but is dispensable for virulence in the corticosteroid murine model of IPA.

**Figure 1.**
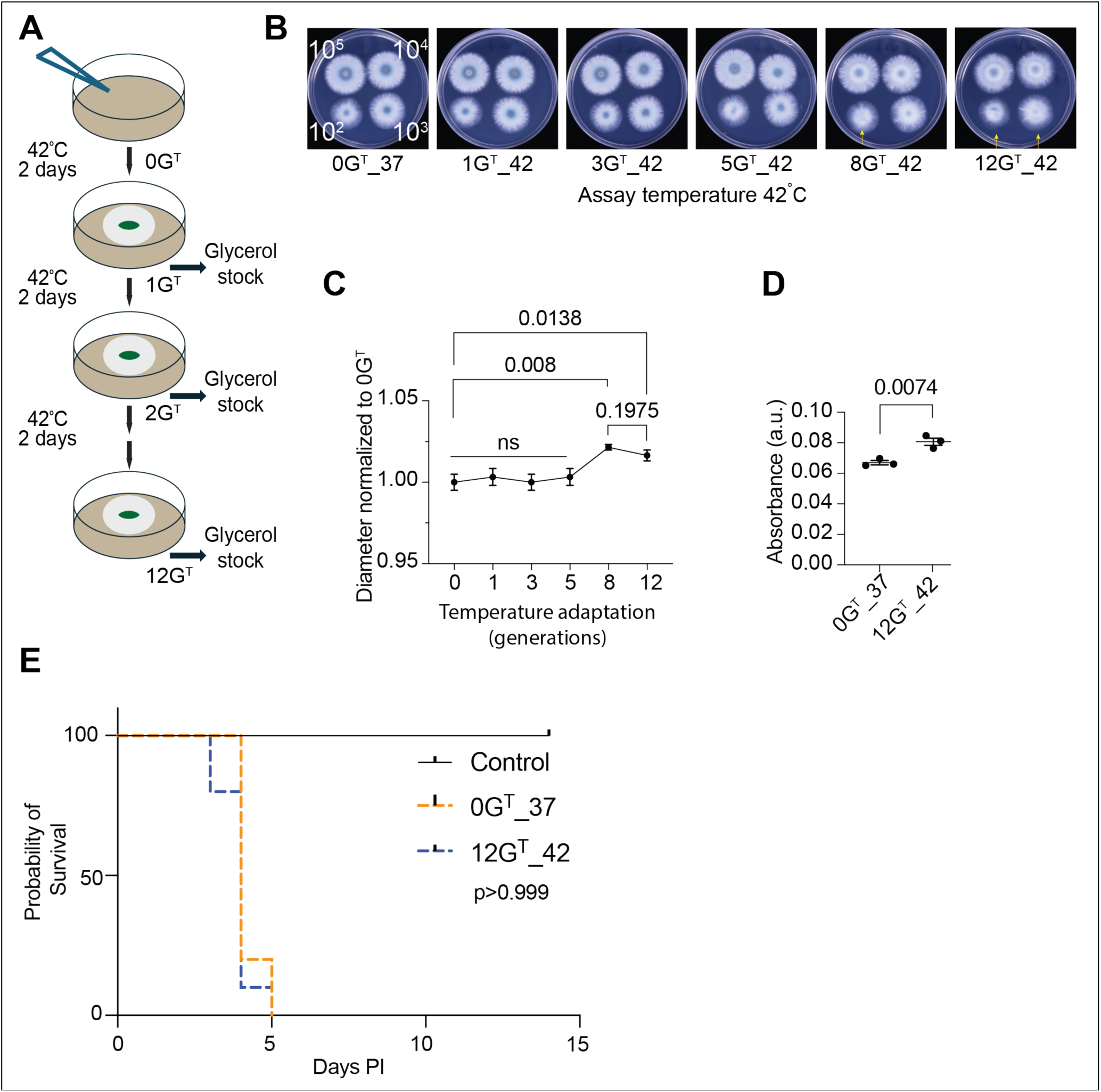
Adaptation of *Aspergillus fumigatus* at 42°C affects morphology and surface attachment. A) Schematic diagram for the temperature adaptation experiment. B) Serial dilutions of indicated generations of the adapted strains grown in GMM plates at 42°C. Different colony morphology observed in 8G^T^ and 12G^T^ strains (yellow arrows). C) Colony diameter normalized to the un-adapted (0G^T^) strain for the colony diameter at 10^5^ conidia. One-Way ANOVA followed by Tukey’s post-hoc analysis was used to compare differences in the mean. p-values are listed for pairwise comparisons. D) Conidia were allowed to adhere to the plastic plate for 12 hours. Biofilms were washed with PBS and GMM was added for additional 12 hours. The Crystal violet assay was used to determine fungal biofilm at 20 hours. One-Way ANOVA followed by Tukey’s post-hoc analysis was used to compare differences in the mean. p-values are listed for pairwise comparisons. Increased surface attachment was observed for the 12G^T^ strain when conidia were collected at either 37°C (12G^T^_37) or 42°C (12G^T^_42). E) Thermal adaptation is dispensable for virulence in a corticosteroid model of IPA. Mice were immunosuppressed and infected with the *A*. *fumigatus* unadapted and adapted strains. Survival was monitored for 14 days. No difference was observed. p>0.999, log-rank test.

### Fungal exposure to increased temperature impacts azole response *in vitro* and *in vivo*

Transcriptional activity before dormancy is linked to stress response, including anti-fungal drug response in *A. fumigatus* (24). Fungal strains exposed to increased temperature in laboratory evolution settings or isolated from compost piles show variable azole response in *Cryptococcus neoformans* and *A. fumigatus,* respectively (17, 19). Thus, to test if temperature-adapted *A. fumigatus* strains have altered response to the azole class of drugs at the human infection temperature, we propagated them at either 37°C (12G^T^_37) or 42°C (12G^T^_42) (Figure 2A). We tested the minimum inhibitory concentration (MIC) for voriconazole and posaconazole using a broth microdilution assay and E-test strips. No change in MIC was observed (Figures 2B and 2C). However, the E-test assay showed a decreased zone of inhibition in the 8G^T^ and 12G^T^ strains and the presence of fungal colonies inside the zone of inhibition (Figure 2C, red congruent circles), indicative of differential azole response as previously shown (22).

**Figure 2.**
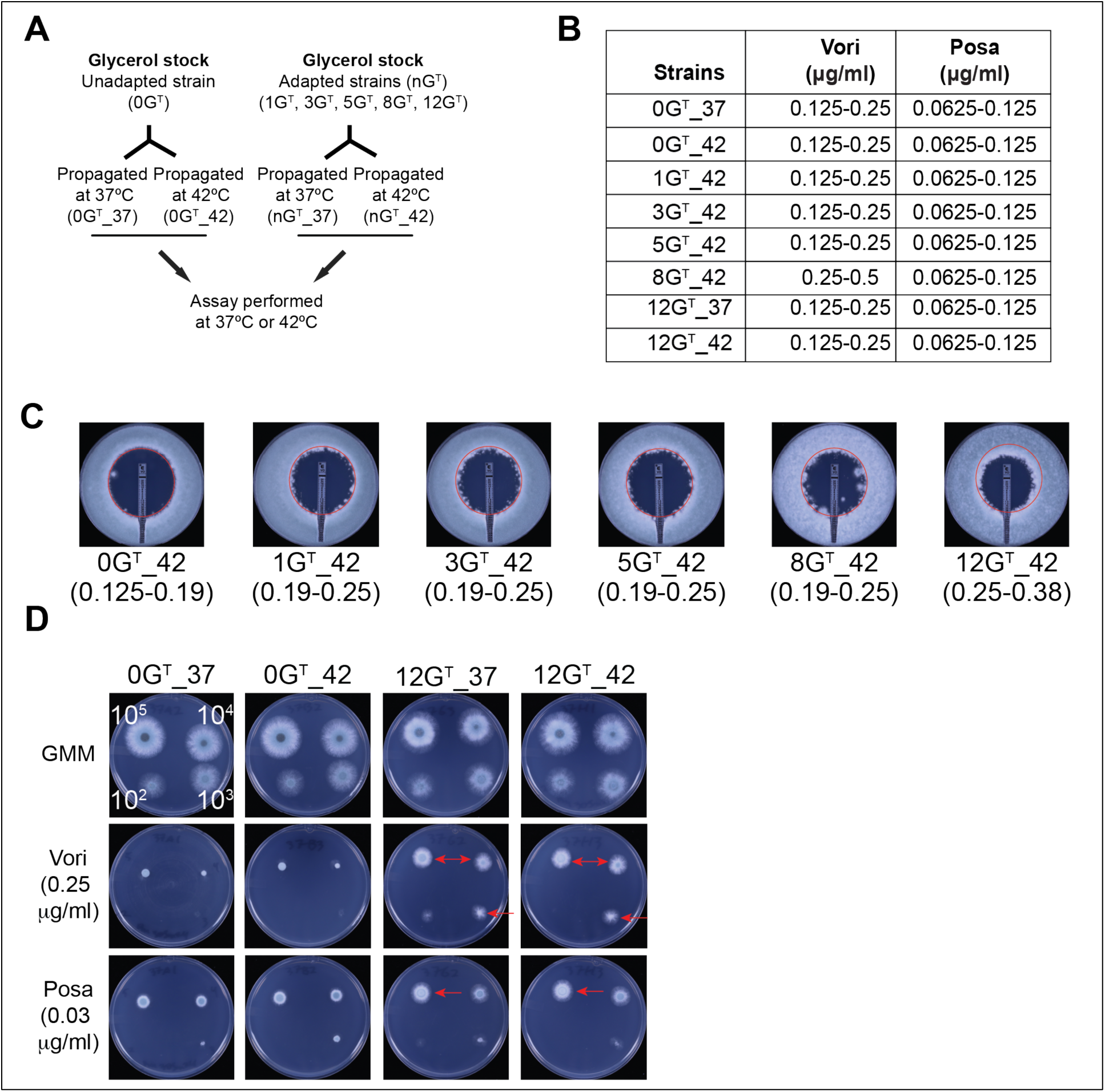
Temperature adaptation crosstalks with azole response in *A*. *fumigatus*. A) Schematic diagram showing the identifiers for unadapted and temperature adapted strains propagated at 37°C or 42°C for the subsequent experiments. B) MIC determination using Broth microdilution assay. MIC values in mg/ml are depicted for voriconazole and posaconazole. No change in MIC was observed. C) E-test assay. MIC was determined using the E-test strip for voriconazole for the indicated strains. The red congruent circle represents the area under the zone of inhibition (ZOI) for 0G^T^. Reduced ZOI was observed upon temperature adaptation. D) Sub-MIC azole response. Serial dilutions of spores collected at either 37°C or 42°C were spotted in the presence of azole drugs as indicated. Red arrows show increased growth in the adapted strain in presence of voriconazole and posaconazole.

To test this response, we plated the WT and 12G^T^ strain on 0.25μg/ml of voriconazole (0.5x MIC) and 0.03μg/ml of posaconazole (0.5x MIC) and observed increased radial fungal growth in the adapted strain at the infection temperature of 37°C irrespective of spores density (Figure 2D, red arrows). Improved fungal growth upon azole exposure is independent of azole target enzyme’s, *cyp51*, transcriptional regulation (Supplementary Figure 2). Previous reports have suggested that CO_2_ potentiates azole activity against *C*. *neoformans* (25). Thus, we tested spore collection and azole response in the presence and absence of supplemented CO_2_. *A*. *fumigatus* isolates grew better in the presence of CO_2_; however, the temperature-azole crosstalk phenotype is independent of CO_2_ (Supplementary Figure 1).

Given the crosstalk between temperature adaptation and azole response, we tested the *in vivo* azole response in a corticosteroid model of *A. fumigatus* infection, as shown in the schematic (Figure 3A) and previously described (22). We observed a significant increase in the fungal DNA in mice infected with the adapted (12G^T^_42) strain compared to fungal DNA from mice infected with the unadapted (0G^T^_37) strain (Figure 3B, p=0.0279, Mann-Whitney U-test). Together, our data showed that temperature adaptation increased fungal growth *in vitro* and increased fungal burden upon treatment with posaconazole in a murine model of infection.

**Figure 3.**
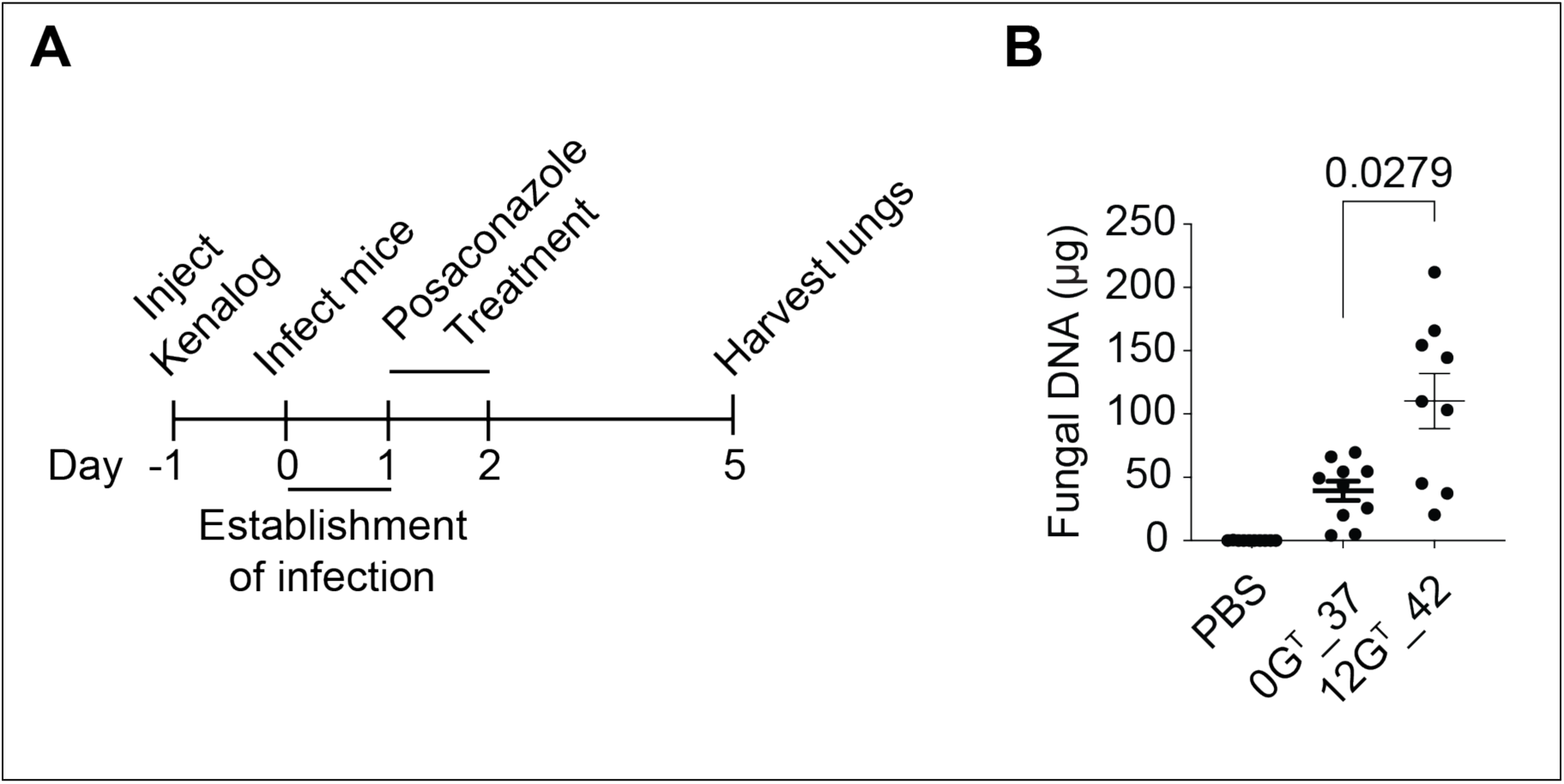
Temperature adaptation increased fungal burden in a murine model of IPA upon azole treatment. A) Schematic diagram showing workflow for harvesting lungs infected with *A. fumigatus* strains in a murine model of IPA. B) Fungal DNA was quantified against a standard curve using a hydrolysis probe and primers specific for fungal 18s rRNA gene. Non-parametric test (Mann-Whitney U-test) was used to determine differences in the mean. p = 0.0279.

### Temperature adaptation is reversible and epigenetic in nature

Our data indicated that temperature selection pressure might be important to maintain adaptation in our adapted 12G^T^ strain. Thus, we hypothesized that this adaptation is reversible. To test this hypothesis, we de-adapted the 12G^T^ strain by repeatedly growing it at 37°C for three generations (12G^T^_RG1 - 12G^T^_RG3, Figure 4A). Notably, the 12G^T^_RG3 strain phenocopies the unadapted strain in presence of azoles. Additionally, the 12G^T^ strain propagated at 37°C still shows improved growth in the presence of azoles, confirming that the phenotype is reversible only after three generations (Figure 4B).

**Figure 4.**
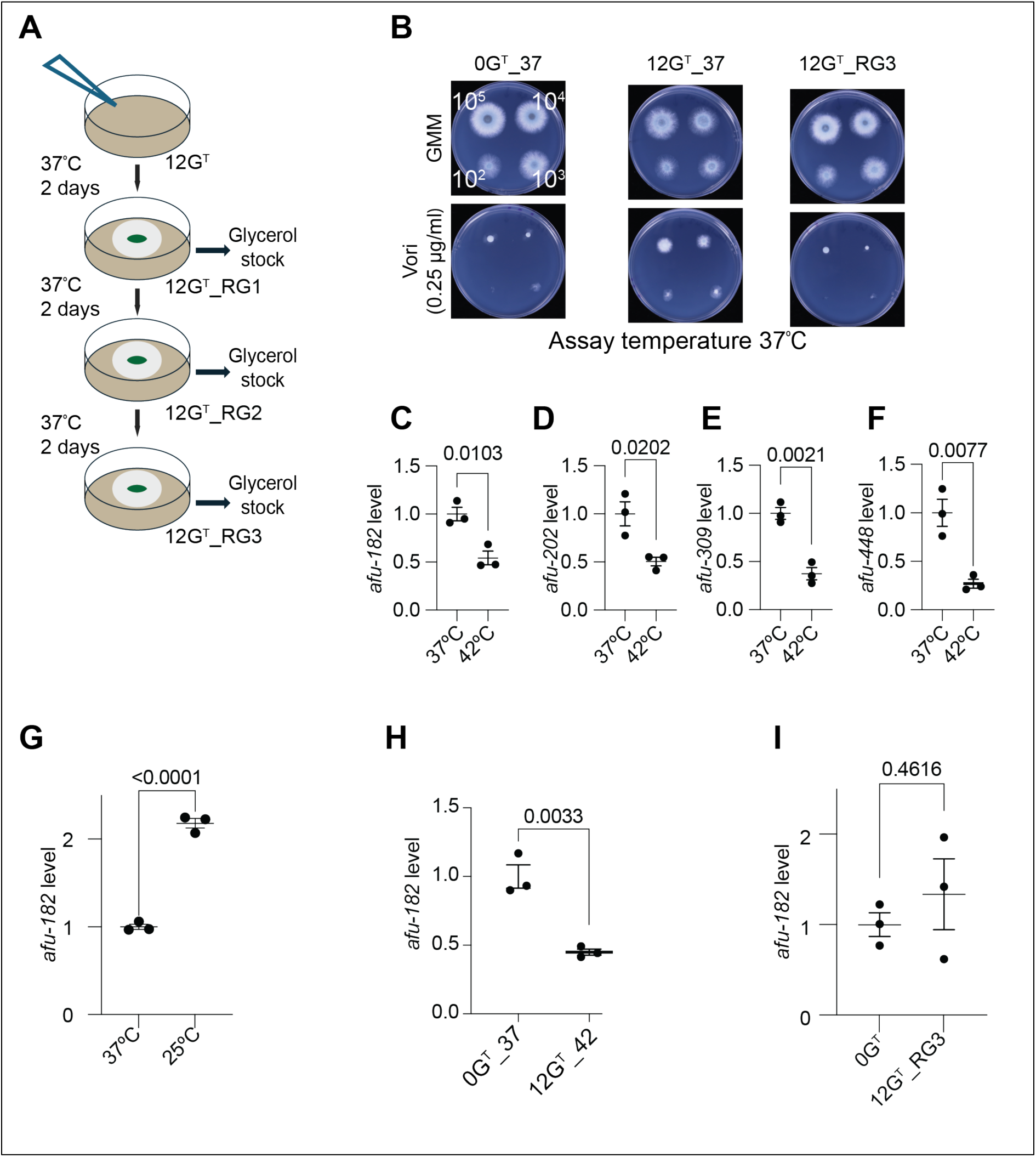
Temperature adaptation is reversible and mediated by lncRNA *afu-182*. A) Schematic diagram for the de-adaptation of temperature adapted strain 12G^T^. B) Fungal conidia of indicated strains were spotted in absence or presence of 0.25μg/ml voriconazole at 37°C. RNA levels of 13 ncRNAs were compared with qPCR at 37°C and 42°C. Four, C) *afu-182*, D) *afu-202*, E) *afu-30S*, and F) *afu-448* showed lower levels at 42°C compared to 37°C. G) RNA levels were quantified at 25°C compared to 37°C. RNA levels of *afu-182* were higher in 25°C compared to 37°C. *afu-182* levels were quantified in H) adapted strain at both 37°C and 42°C compared to unadapted strains or I) in the 12G^T^_RG3 strains. Lower *afu-182* levels were observed in the 12G^T^ adapted strain and *afu-182* levels revert to WT levels in the 12G^T^_RG3 strain. Two-tailed student’s t-test was used to compare means between two groups (C-G and I), and One-Way ANOVA followed by Tukey’s post-hoc analysis was used to compare differences in the means in H.

Epigenetic and reversible mechanisms behind altered antifungal responses have been previously reported in fungi (26, 27). We have previously shown that a lncRNA, *afu-182*, mediates sub-MIC response in *A. fumigatus* (22). We therefore tested the role of lncRNAs in temperature adaptation. We assessed the RNA levels of 13 previously known ncRNAs (28) in *A. fumigatus* grown at 42°C compared to 37°C. Four RNAs, *afu-182* (p=0.0103), *afu-202* (p=0.0202), *afu-30S* (p=0.0021), and *afu-448* (p=0.0077) showed significantly lower levels at 42°C compared to 37°C (Figures 4C-F and Supplementary figures 3A-I, student’s t-test).

To confirm if the RNAs showing differential response are temperature regulated RNAs, we compared the RNA levels of these 4 ncRNAs at 25°C compared to 37°C; only *afu-182* showed increased levels at 25°C (Figure 4G, p<0.0001, student’s t-test and supplementary figures 3J-L). Importantly, the colony growth at 25°C is much smaller than at 37°C and 42°C. Thus, to determine if colony size affected the *afu-182* RNA levels, we collected the RNA from *A. fumigatus* colonies at 37°C at different growth stages. No difference in *afu-182* levels was observed (Supplementary 4), indicating that *afu-182* levels are responsive to temperature and not colony diameter. Thus, *afu-182* RNA levels negatively correlate with increasing temperature.

### Thermal adaptation and azole response crosstalk is mediated by lncRNA *afu-182*

As the temperature adaptation phenotype is reversible, we tested the RNA levels of *afu-182* in 12G^T^ and 12G^T^_RG3 strains compared to the unadapted (0G^T^) strain. Interestingly, we see a significant reduction in *afu-182* RNA levels in the 12G^T^ strain (Figure 4H, p=0.0033, student’s t-test), and *afu-182* levels revert to WT levels in the deadapted strain,12G^T^_RG3 (Figure 4I, p=0.4616, student’s t-test). To determine if *afu-182* levels explain the azole phenotype in 12G^T^ strains, we ectopically expressed *afu-182* driven by *gpdA* promoter (referred to as 12G^T^_P, P indicating plasmid) as previously described (22) at 42°C. The resultant strain 12G^T^_P strain phenocopies the azole stress response of the unadapted strain at 42°C (Figure 5A) and has *afu-182* levels similar to the unadapted strain (Figure 5B, p = 0.5235, student’s t-test), showing that RNA levels of *afu-182* mediate temperature and azole crosstalk. Reverting the temperature adaptation or ectopically increasing the *afu-182* levels is sufficient to revert the azole susceptibility of the temperature adapted strain.

**Figure 5.**
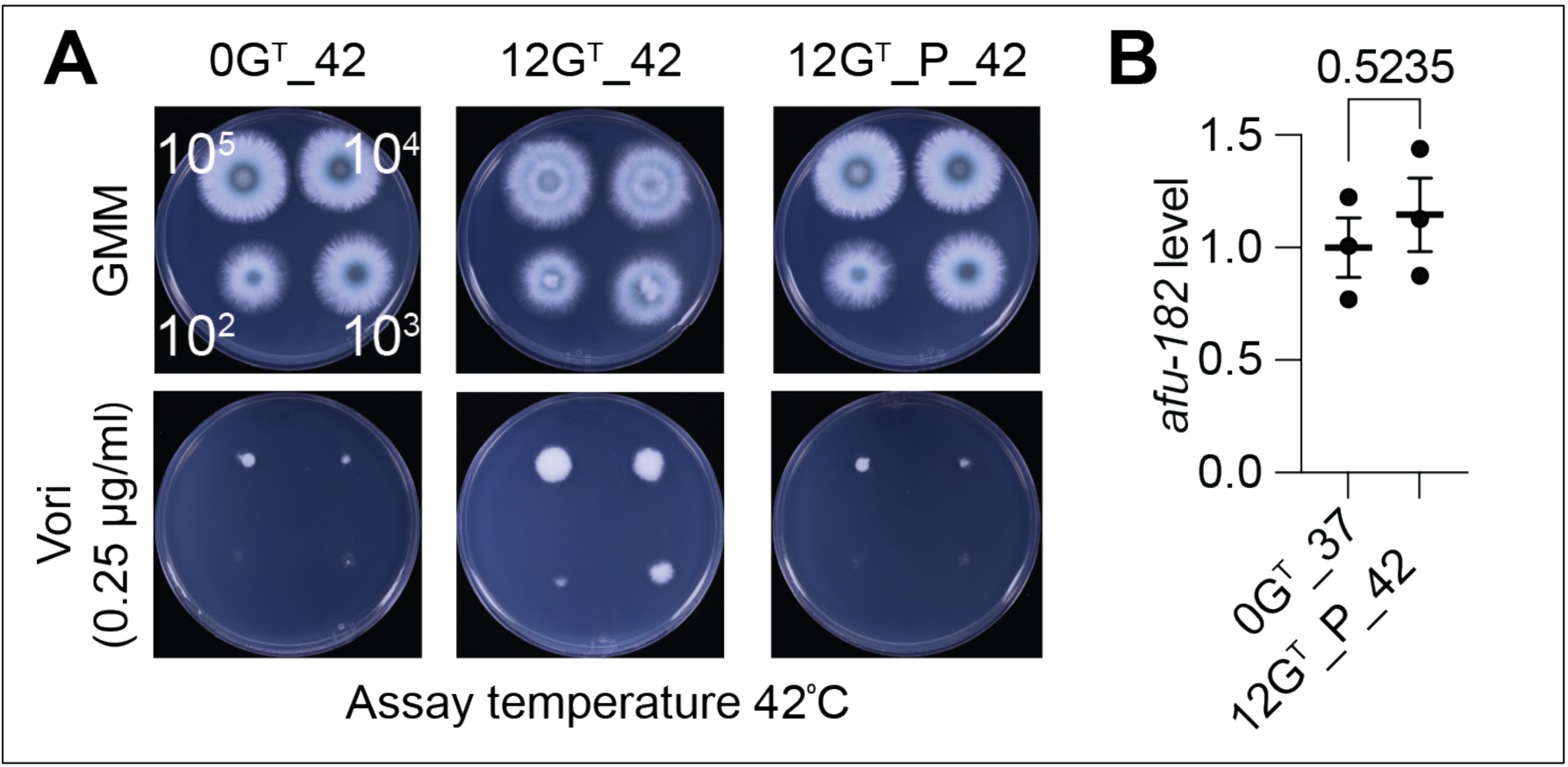
Ectopic expression of afu-182 at 42°C reverses the azole phenotype in temperature adapted strains. A) Spores from indicated strains were spotted as indicated in the absence or presence of 0.25μg/ml voriconazole. B) RNA levels of *afu-182* were quantified using qPCR. Two-tailed student’s t-test was used to compare means between the groups.

### Overexpression of *afu-182* improves outcomes in infections caused by azole-resistant isolates

We previously demonstrated that *afu-182* is a negative regulator of azole drug response, both *in vitro* and *in vivo* (22). Deletion of *afu-182* increased fungal growth in the presence of sub-MIC azole concentrations in a corticosteroid murine model of IPA, and overexpression of *afu-182* in CEA10 decreased fungal growth *in vivo* (22). Here, we over-expressed the lncRNA *afu-182* in the azole-resistant strains 08-19-02-10 (29) and F262del (30) (Supplemental figure 5A and 5B). Over-expression of *afu-182* did not change the MIC in either strain background (Figure 6A and 6B) but showed increased inhibition in fungal growth in sub-MIC concentrations (Figure 6C and 6D). To study the role of this phenomenon in azole treatment outcomes in drug resistant isolates, we used a corticosteroid model of IPA as previously described (22). Mice infected with the 08-19-02-10 host strain, and the 08-19-02-10 OE-182 strain were treated with 20 mg/kg posaconazole (Figure 5E), and infected with F262del host and F262del OE-182 strain were treated with 5 mg/kg (Figure 5G) based on the MIC of the strains for posaconazole. Overexpression of lncRNA *afu-182* in both strain backgrounds led to significant survival improvement (Figures 5F and 5H, p=0.0.0361 and p=0.0.0148, respectively, log-rank test) compared to the host strains even in the absence of a change in MIC. No significant differences in virulence were observed due to OE *afu-182* in the absence of azole treatment (Supplementary Figure 5C and 5D).

**Figure 6.**
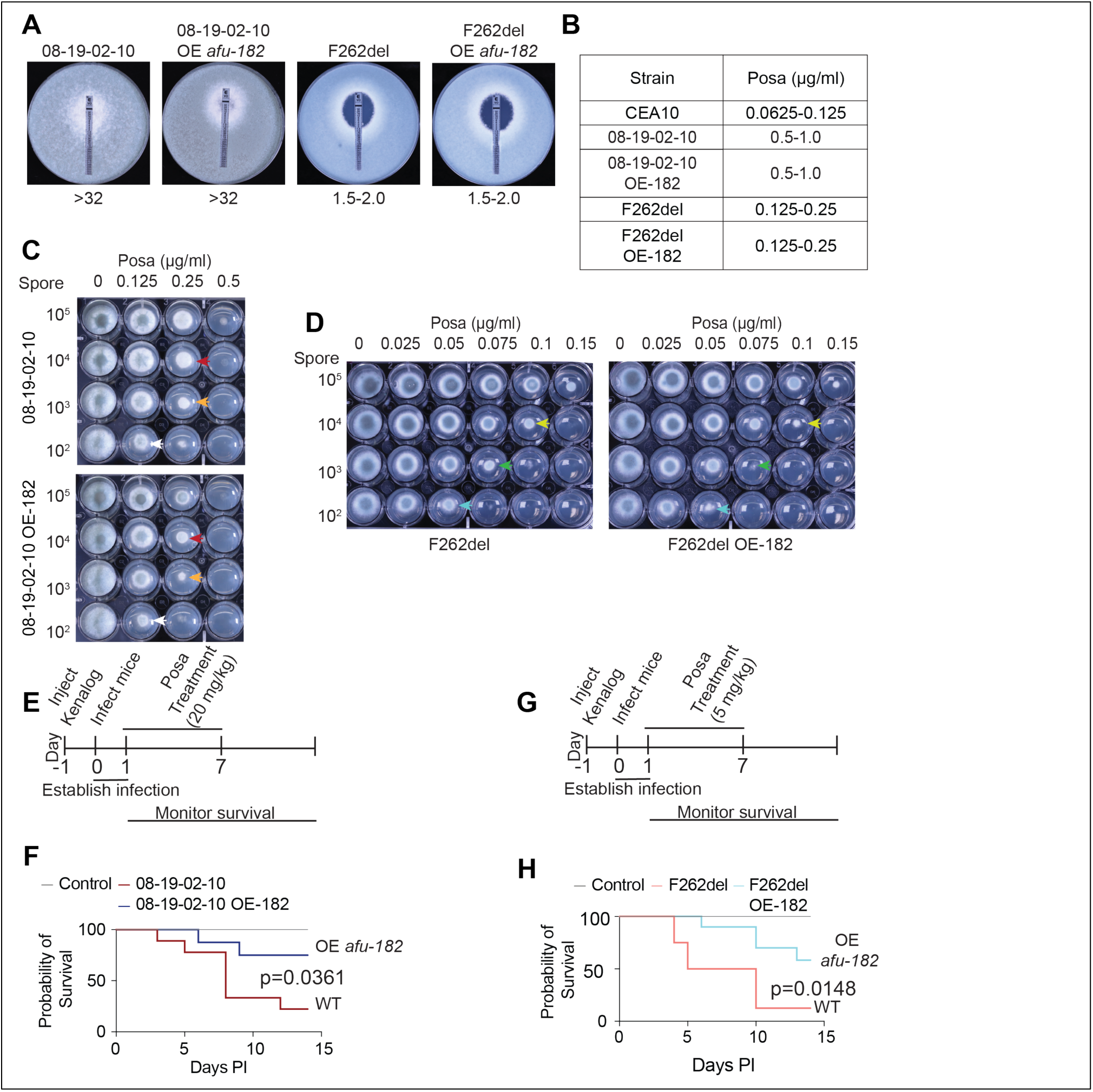
Overexpression of *afu-182* increases survival in a murine model of IPA infected with azole resistant *A. fumigatus* isolates. MIC values as determined by A) E-test assay. or B) broth microdilution assay for the indicated strains. Spot assay for C) Strains 08-19-02-10 and 08-19-02-10 OE-182, and D) for Strains F262del and F262del OE-182 at the indicated drug concentration. Same color arrows mark the wells with the same spore and drug concentrations, showing the effect of sub-MIC concentration of posaconazole on OE *afu-182* strains. Schematic diagrams (E and G) showing workflow for survival analysis upon azole treatment at the indicated concentrations. Survival analysis for mice infected with F) strains 08-19-02-10 and 08-19-02-10 OE-182, and H) F262del and F262del OE-182. Log-rank test was used to assess the survival difference between the strains.

## Discussion

Environmental *A. fumigatus* strains differ in their virulence potential (31); however, the true origin of environmental samples is up for debate, as an environmental *A. fumigatus* strain may adapt in animals or birds (37°C-43°C) or compost before re-entering as “environmental” samples. It is evident that fungi can adapt to different temperatures, and pathogenic fungi are good colonizers of the mammalian lung (32). However, how prior temperature adaptation affects virulence, including azole drug response, is not completely understood. We found that *A. fumigatus* adaptation to increased temperature led to increases in colony diameter and surface attachment that may have virulence significance (Figures 1C and 1D). Although the thermal adaptation was dispensable for virulence (Figure 1E), previous reports have shown a correlation between temperature adaptation and azole resistance in the pathogenic yeast *C*. *neoformans*(19). Thus, we tested the azole response of the 12G^T^ strain and observed that thermal adaptation did not change the azole MIC (Figures 2B and 2C); however, it resulted in increased colony growth in the presence of sub-MIC azole concentrations (Figure 2D), indicating the crosstalk between temperature and azole response in *A. fumigatus*. To ascertain if this crosstalk affects azole treatment outcomes, we tested the azole response phenotype in a murine model of IPA, and the 12G^T^ strain resulted in increased fungal burden in murine lungs upon azole treatment (Figure 3B). It is possible that a selection pressure of either increased temperature or azole stress response is important to maintain this adaptation, and thus a crosstalk between these two stresses. We de-adapted our 12G^T^ strain at 37°C for three generations (12G^T^_RG3), resulting in a reversion to the WT phenotype (Figure 4B) and confirming the transient nature of this phenotype. It is plausible that due to hypothermia (<29°C) experienced by mice during *A*. *fumigatus* infection (33), the adaptation reverts in vivo, leading to no difference in survival compared to unadapated strain (Figure 1E). As we cannot rule out the role of hypothermia in reversing the phenotype of the 12G^T^ strain in the murine model of IPA, thermal adaptation may increase fungal virulence in the absence of azole treatment in other animal models or in humans experiencing febrile temperatures during infection.

Epigenetic mechanisms involving RNAi for transient drug resistance have been described previously in *Mucor circenolloides* (34). We have previously shown that lncRNA *afu-182* regulates fungal response to sub-MIC azole drugs; thus, we tested the RNA levels of 13 previously identified ncRNAs (28). Interestingly, we observed 4 ncRNAs to have a negative correlation between 37°C and 42°C (Figures 4C-F). However, we observed only lncRNA *afu-182* levels to be negatively regulated by temperature at 25°C (Figure 4G and supplementary figures 3J-L). To understand the role of afu-182 in temperature-azole crosstalk, we ectopically expressed the *afu-182* while keeping the *A. fumigatus* 12G^T^ strain constantly at 42°C to prevent temperature mediated changes. Increasing the *afu-182* levels back to WT ectopically (Figure 5B) leads to loss of fungal sub-MIC azole growth (Figure 5A), indicating that *afu-182* mediates the crosstalk. Importantly, during the course of this study, additional ncRNAs in *A*. *fumigatus* have been identified (35). If other ncRNA mediated mechanisms play a role in this crosstalk, it is currently unknown.

The first report of azoles’ anti-fungal property was presented in 1944 (36), highlighting the question of *afu-182’s* ecological presence in *A*. *fumigatus*. We have previously shown that *afu-182* only affects fungal response to azoles and not to polyenes and echinocandins (22). It is possible that *afu-182’s* major role is in fungal temperature adaptation. As ncRNAs have emerged as a regulator of stress response, this reversible ability possibly enables *A*. *fumigatus* to adapt between saprophytic and pathogenic lifestyles. This hypothesis correlates with the increased isolation of ARAF strains from soil with compost that may be exposed to high temperatures (17).

The mechanisms of *afu-182-*mediated azole response are still under investigation but appear to be independent of *cyp51* transcriptional regulation (Supplementary Figure 2). Further studies comparing fungal response to temperature and azoles may shed more light on the molecular aspects of this crosstalk. Importantly, azole exposure is considered the primary route of azole resistance; however, we previously showed that in a laboratory adaptation experiment to low dose azole, the change in MIC is *afu-182* dependent (22). On the other hand, and more importantly, over-expression of *afu-182* in clinical drug-resistant isolates with varying resistance mechanisms improves disease outcomes in a corticosteroid model of IPA (Figures 6F and 6H). In both cases, azole MIC values do not change, and overexpression of *afu-182* does not affect virulence in the absence of azole treatment (Supplementary figures 5C and 5D).

As MIC values did not change *in vitro* but improved disease outcomes in ARAF, it is possible that multiple stresses in the animal model, along with azole drugs, impact *in vivo* fungal growth. It is also possible that the drug concentration at the site of infection is below the MIC values, thus showing the same effects as observed *in vitro*. It is known that the drug concentration at the infection site is lower than the serum concentration (37), the effect of this variability is currently not known in IPA treatment. Thus, a more thorough understanding of fungal azole response is needed before classification as a susceptible or resistant isolate, especially when deciding on a treatment regimen.

Taken together, our data show *A. fumigatus* temperature adaptation is associated with azole response and is mediated by lncRNA *afu-182*. However, how *afu-182* functions is currently unknown. The structures and functions of lncRNAs have been studied in various organisms (38–40), however, their mechanistic characterization is still lacking in *A. fumigatus*.

Structural analysis of *afu-182* at different temperatures and determination of its binding partners will highlight the functions of *afu-182* and bridge the knowledge gap of azole response and temperature regulation. Given that the OE-182 strain improves disease outcomes in ARAF strains with differing resistance mechanisms, further mechanistic analysis of *afu-182* interacting partners allows an avenue for developing novel therapies for drug resistant isolates. As we have shown previously, low dose exposure to azoles leads to an increase in MIC only in Δ*afu-182* strain (22), studying the *afu-182* interactome will provide a framework to prevent resistance from arising, especially *in vivo*.

## Materials and methods

### Fungal strains

*Aspergillus fumigatus* strains used in the study are listed in Table 1. All strains were stored as glycerol stocks at −70°C for long term storage. Strains were propagated in glucose minimal media (41) at 37°C or 42°C as indicated. Agar (1.5%) was added to the media for plating. The adaptation experiments were carried out in the absence of CO_2_. For all other experiments, strains were grown in the presence of 5% CO_2_ to mimic CO_2_ stress during infection unless stated otherwise. The strains and isolates of *A. fumigatus* used in this study are listed in Table 1.

**Table 1:** List of strains used in this study.

| S.N. | Strains | Names used in the paper | Citation |
| --- | --- | --- | --- |
| 1 | CEA10 (Unadapted) | CEA10 or Unadapted or 0G<sup>T</sup> | Cramer Lab |
| 2 | CEA10_ Temperature adapted for 1 generation | 1G<sup>T</sup> | This study |
| 3 | CEA10_ Temperature adapted for 3 generations (Gen3) | 3G<sup>T</sup> | This study |
| 4 | CEA10_ Temperature adapted for 5 generations (Gen5) | 5G<sup>T</sup> | This study |
| 5 | CEA10_ Temperature adapted for 8 generations (Gen8) | 8G<sup>T</sup> | This study |
| 6 | CEA10_ Temperature adapted for 12 generations (Gen12) | 12G<sup>T</sup> | This study |
| 7 | DeAdapted Gen12 for three generations at 37°C (12GT_RG3) | 12G<sup>T</sup>_RG3 | This study |
| 8 | Gen12 strain transformed with plasmid for Overexpression of *afu-182* (12GT_P) | 12G<sup>T</sup>_P | This study |
| 9 | *afu-182* deletion $\Delta afu-182$ | $\Delta afu-182$ | (22) |
| 10 | 08-19-02-10 | 08-19-02-10 | (43) |
| 11 | 08-19-02-10 OE *afu-182* | 08-19-02-10 OE *afu-182* | This study |
| 12 | CEA10 *hmg1 F262del* | F262del | (44) |
| 13 | CEA10 *hmg1 F262del* OE *afu-182* | F262del OE *afu-182* | This study |

**Table 2:**
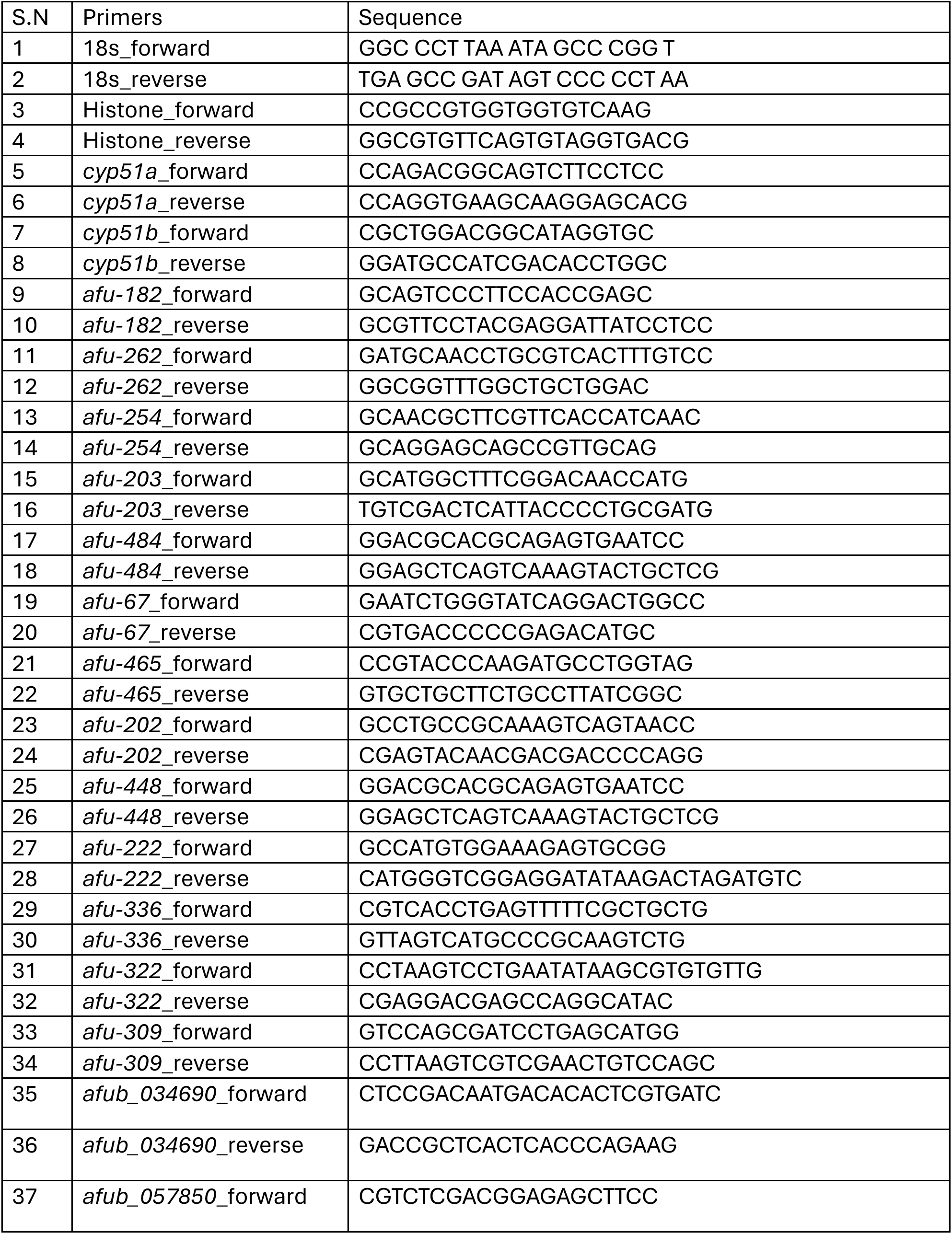
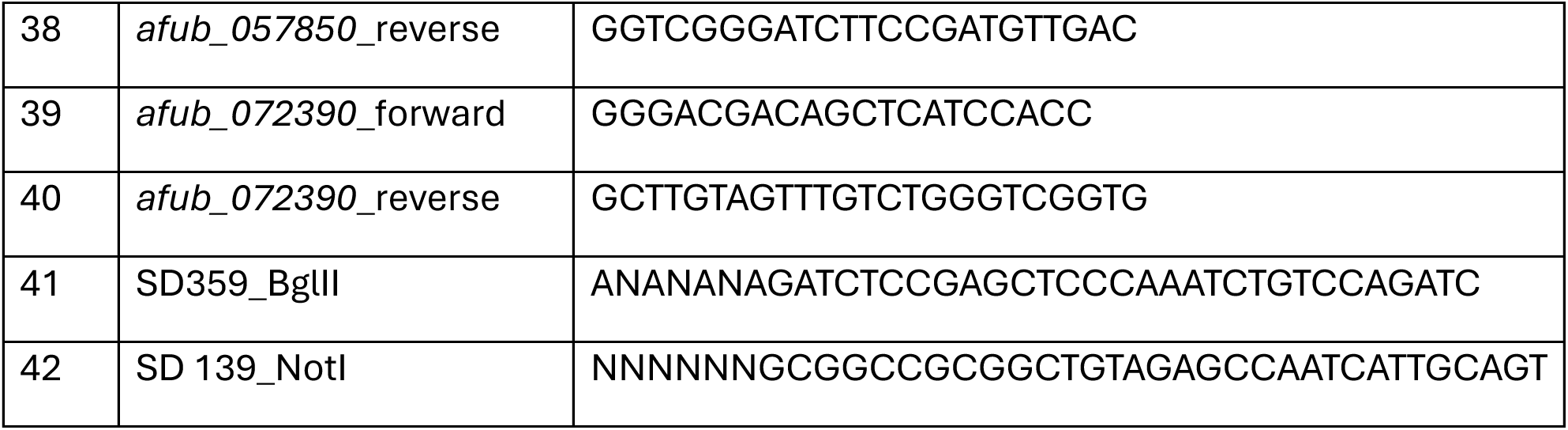
List of primers used in this study.

### Adaptation experiment

*A. fumigatus* strain CEA10 was adapted at 42°C for 12 generations (Figure 1A). Briefly, CEA10 unadapted (0G^T^) strains were spot inoculated in a glucose minimal media plate and incubated at 42°C without CO_2_ for 48 hours. Spores from the first generation of temperature adapted strain (1G^T^) were collected in 0.01% Tween-80, and again spot inoculated onto a fresh GMM plate for adaptation to the next generation. The process was repeated until spores for the 12^th^ generation of temperature adapted strains were collected and stored as glycerol stock. Spores were propagated at either 37°C or 42°C for the subsequent experiments as indicated. The naming scheme is shown in the schematic (Figures 1A and 2A). The temperature adapted strains were propagated in presence of 5% CO_2_ unless stated otherwise.

#### OE afu-182ptrA plasmid construction

*A. nidulans gpdA*(p)::afu-182 fragment was amplified from OEafu-182pyrG plasmid (22) using primers SD359_BglII and SD 139_NotI. The fragment was digested with enzymes BglII and Not1 and was ligated into plasmid pTDS8 previously digested with BglII and Not1 resulting in plasmid OEafu-182 ptrA.

### Generation of OE-182 strains

OE-182 strains were generated as described previously (22). Briefly, host strains (12G^T^, 08-19-02-10, and Fdel262) were transformed with OE-182 ptrA plasmid for ectopic insertion using polyethylene glycol mediated DNA transformation of fungal protoplast (22). All strains were confirmed for the presence of the plasmid with PCR. RNA levels were confirmed by qPCR using ssoA Advanced SYBR mix (BioRad) as described. Multiple transformants for each host strain were selected and confirmed.

### Spot assay

Plates containing solid GMM were spot inoculated with *A. fumigatus* spores (1x 10^5^, 1x 10^4^, 1x 10^3^, 1x 10^2^) and grown at the indicated temperatures and indicated concentration of drugs. For assays performed in 24-well plates, 1 ml of media was added to each well, and the indicated spore concentration was inoculated. All experiments were done in triplicate and were repeated three times.

### Biofilm formation assay

*A. fumigatus* spores (1×10^5^/ml) in liquid GMM were inoculated in a 12-well plate. The plates were centrifuged at 250g for 10 min and incubated at 37°C for 12 hours. The crystal violet stain was performed after washing the biofilms twice with PBS as described previously (22). Absorbance was measured using a plate reader (Multisan SkyHigh, Thermo scientific) at 600 nm. All experiments were done in triplicate, and three independent experiments were performed.

### E-test assay

Five milliliters of top agar (0.75%) containing 1×10^5^ *A. fumigatus* spores was gently spread on GMM plates and allowed to solidify. An E-test strip (bioMérieux) was carefully laid on the top of the plate per manufacturer’s recommendation. The plates were incubated at 37°C for 48 hours, and the minimum inhibitory concentration (MIC) was recorded. The experiment was done in triplicate for each strain and repeated three times.

### Broth Microdilution Assay

Azole drugs were column-wise serially diluted (1:2) in 100ml of GMM. Then, 100ml of *A. fumigatus* spores (5×10^3^) in GMM were added. The visible growth of *A. fumigatus* strains was observed in each of the wells, and the lowest concentration of azoles showing no growth for an *A. fumigatus* strain was recorded as MIC. The experiment was done in duplicate and repeated three times.

### Animal Survival analysis and fungal burden

Outbred CD-1 mice (Charles River Laboratories) weighing 18-23g were housed (5 per cage) in micro-isolator cages. The mice were kept in a soft-lit, well-ventilated cubicle with a 12h:12h light/dark cycle. Mice were immunosuppressed with a single subcutaneous injection of 40 mg/kg body weight of Triamcinolone acetonide (Kenalog-10; Bristol-Myers Squib) on day −1. Mice were briefly anesthetized with isoflurane and infected by intranasal instillation of 2’10^6^ spores in 40 µl PBS. Ten mice were infected per *A. fumigatus* strain for survival analyses, whereas five mice per strain were used for fungal burden analyses. The mice were observed for 14 days for their survival. For azole treatment, Posaconazole (Noxafil, Merck) was diluted in PBS and given via oral gavage at the indicated concentrations for 7 days starting after 24 hours post-inoculation.

For fungal burden, the mice were treated twice with 5 mg/kg posaconazole after 24 and 48 hours of spore inoculation, and the mouse lungs were collected on day 5. The mouse lungs were lyophilized overnight, homogenized with 2.3 zirconia beads, and lysed with LETS buffer (0.1M LiCl, 10mM EDTA pH 8.0, 10mM Tris-Cl pH 8.0, 0.5% SDS). Total DNA was extracted in an aqueous phase using phenol: chloroform: isoamyl alcohol and precipitated with isopropanol. Fungal burden was determined by calculating the levels of 18s rRNA encoding gene against a standard curve using iTaq universal probe mix and hydrolysis probe as previously described (22).

### RNA extraction and assessment of gene expression by RT-qPCR

For RNA sequencing, total RNA was extracted from *A. fumigatus* strain CEA10 (1 x 10^4^ spores) grown at 25°C, 37°C, and 42°C for 72 hours in solid GMM plates. The samples were homogenized with 2.3 mm zirconia beads in Acid Guanidinium Thiocyanate Phenol chloroform (AGPC) (42), treated with Turbo DNAse (Thermo Fisher) per manufacturer’s recommendation and purified using magnetic beads (Beckman Coulter)

For qPCR, 48-hour-old *A. fumigatus* colonies grown in solid GMM plates were homogenized in AGPC, treated with turbo DNAse per the manufacturer’s recommendation. First strand cDNA was synthesized using 500 ng of DNAse treated RNA using MMLV reverse transcriptase (Promega) per the manufacturer’s recommendation. Quantitative PCR (CFX real-time PCR system, Bio-Rad) was done using the sso Advanced universal SYBR green supermix (Bio-Rad). Histone H4 was used as a reference gene. All primers used in the study are listed in Supplementary Table 2.

### Statistical Analysis

GraphPad Prism version 10 was used for all statistical analyses. Parametric test (Student’s t-test or One-Way ANOVA or Two-Way ANOVA followed by Tukey’s post-hoc analysis) was used for in vitro experiments. Non-parametric test (log-rank for survival and Mann-Whitney U test) was performed for animal data. Exact p-values are represented on graphs. All in vitro experiments were performed in biological triplicate and were repeated three times.

## Acknowledgements

We would like to acknowledge the Eukaryotic Pathogen Innovation Center for discussions surrounding the experiments in the manuscript. The research in the Dhingra lab is supported by start-up funds from the Department of Biological Sciences at Clemson University and a National Institute of General Medical Sciences (NIGMS) award (P20GM146584) (PI - James Morris). The funding sources had no role in the study design, data collection and interpretation, preparation of this manuscript or the decision to submit the manuscript. We thank Dr. Jarrod Fortwendel (UTHSC) for Fdel262 strain.

---

**Supplementary Figure 1.**
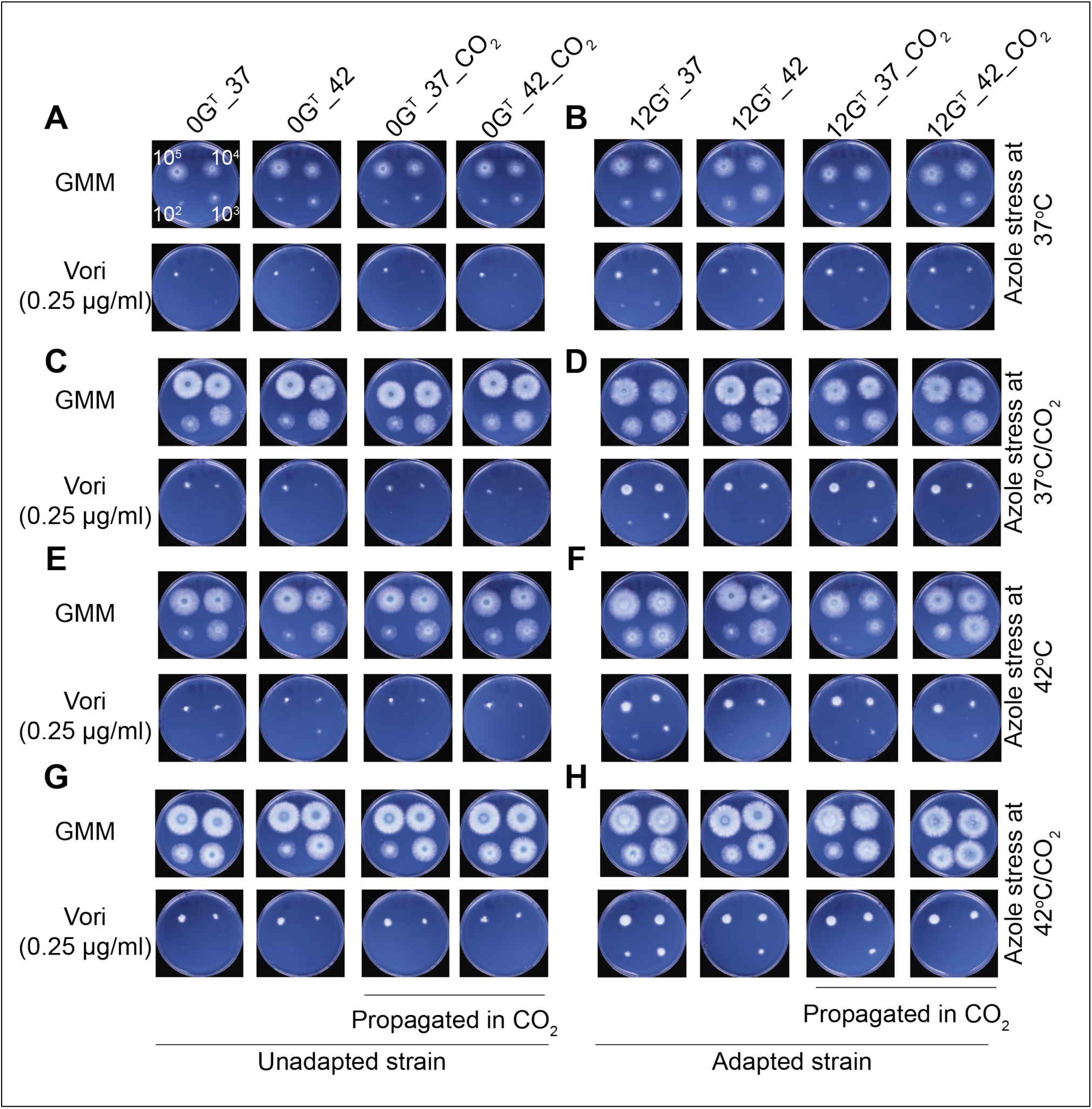
Temperature-azole crosstalk is independent of CO_2_. Unadapted strain 0G^T^ (A, C, E and G) and temperature adapted strain 12G^T^ (B, D, F, H) were propagated at 37°C (labeled as - _37) or 42°C (labeled as - _42) without or with CO_2_ (labeled as - _CO_2_) and were grown with or without voriconazole (0.25μg/ml) in absence or presence of CO_2_. No difference due to CO_2_ was observed.

**Supplementary Figure 2.**
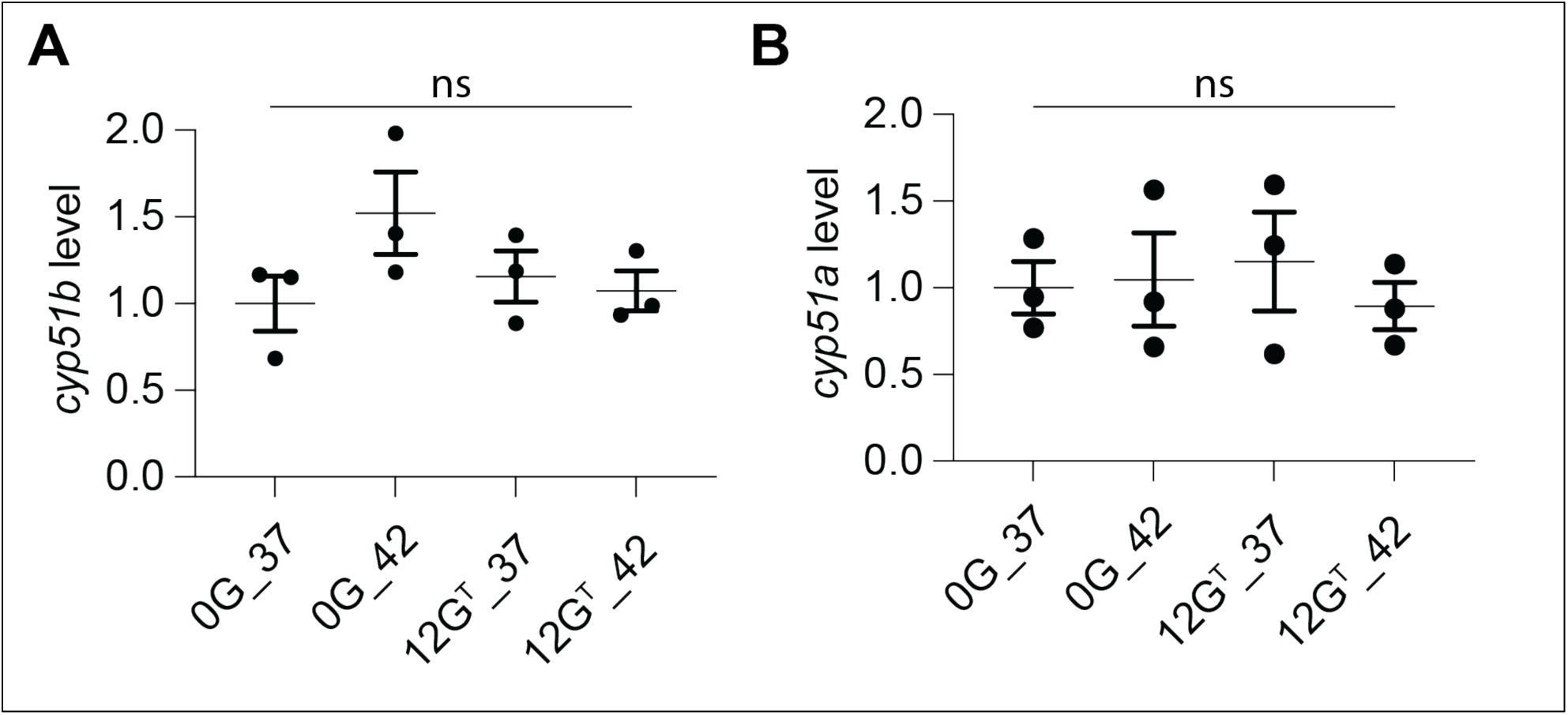
Temperature-azole crosstalk is independent of *cyp51* transcriptional regulation. RNA was extracted from samples either propagated at 37°C or 42°C from colonies grown on GMM plates. No difference in mRNA levels was observed upon temperature adaptation.

**Supplementary Figure 3.**
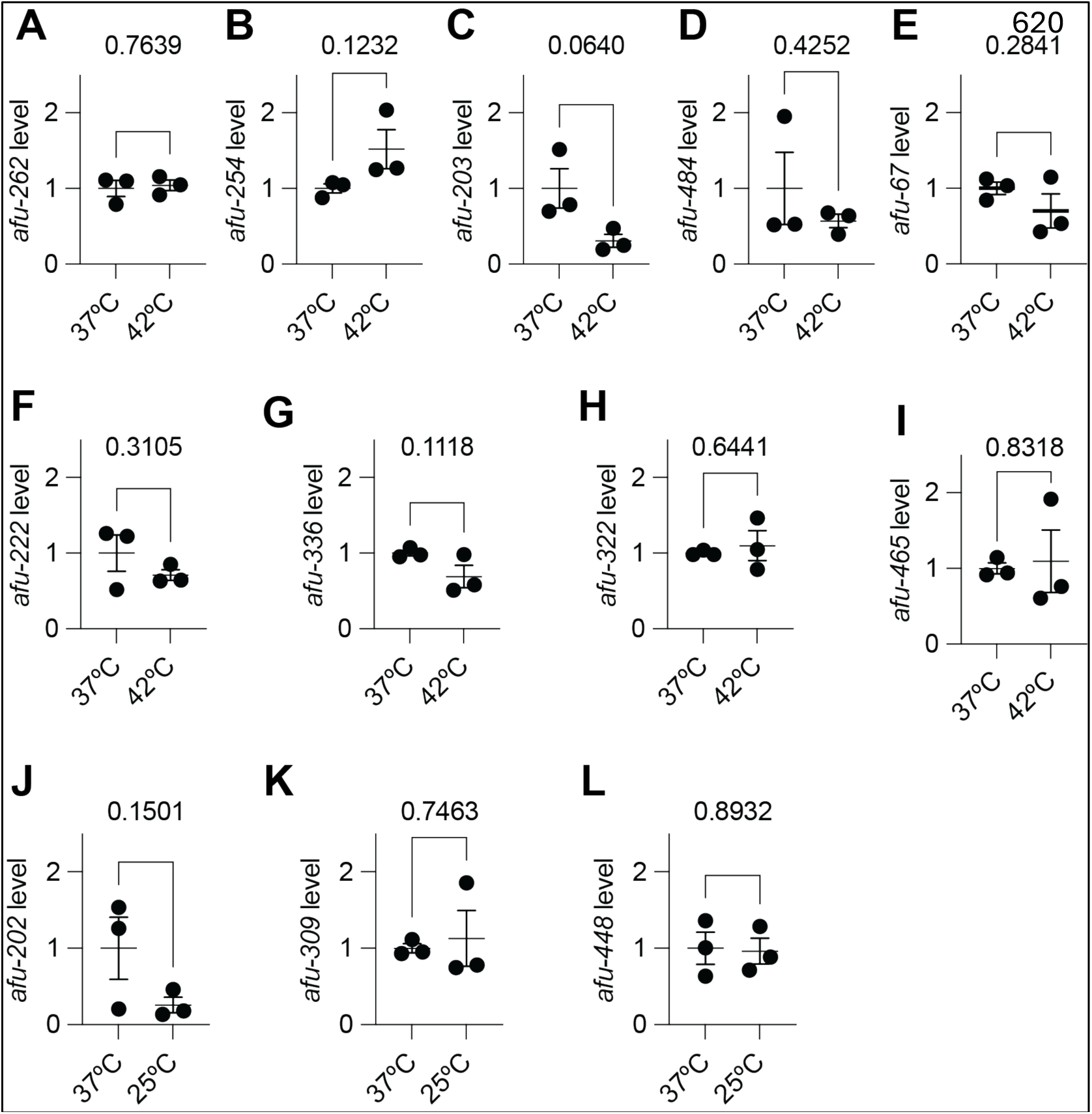
RNA levels of other non-coding RNA levels at 42°C and 25°C. (A-I) RNA levels of ncRNAs were quantified at 37°C and 42°C. (J-L) RNA levels were quantified for RNA showing difference for J) afu-202, K) afu-309 and L) afu-448 at 25°C. No difference was observed at 25°C. Exact p-values are represented (two-tailed Student’s t-test)

**Supplementary Figure 4.**
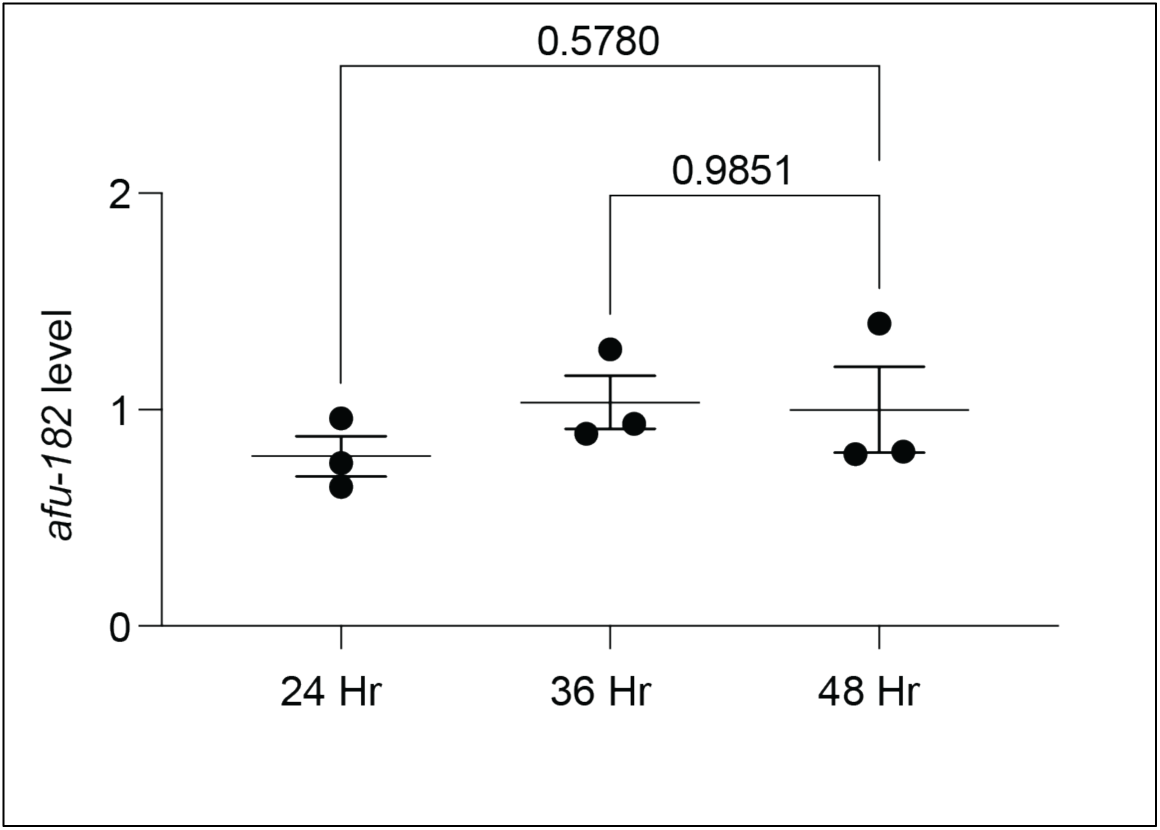
*afu-182* RNA levels are independent of colony size. *afu-182* RNA levels were measured at different indicated time points. No significant difference was observed (One-Way ANOVA followed by Tukey’s pos-hoc analysis).

**Supplementary Figure 5.**
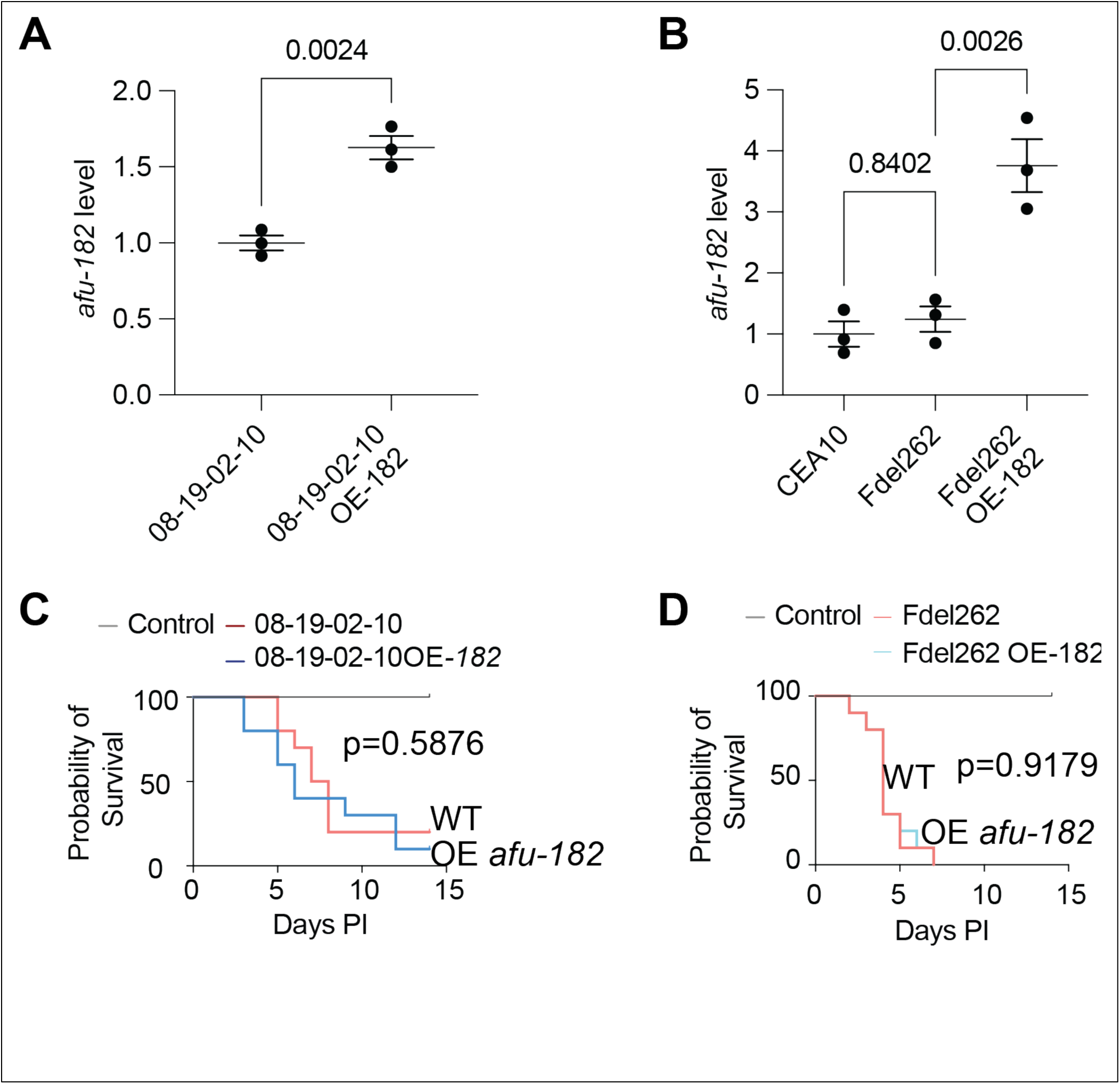
Overexpression of *afu-182* is dispensable for virulence in the corticosteroid murine model of infection. *afu-182* RNA levels were quantified in the OE *afu-182* strain in A) 08-19-02-10 (two-tailed student’s t-test) and B) F262del host strain (CEA10 background, One-Way ANOVA followed by Tukey’s post-hoc analysis). Survival assay for mice infected with strains in C) 08-19-02-10 and 08-19-02-10 OE afu-182 and D) F262del and F262del OE *afu-182* backgrounds. No significance in virulence was observed (Log-rank test).

## Notes

### Competing Interest Statement

The authors have declared no competing interest.

### Summary of Updates

The manuscript is shortened. Figure 7 and Supplementary figure 6 have been removed.

